# Cell divisions both challenge and refine tissue boundaries in the *Drosophila* embryo

**DOI:** 10.1101/2025.03.31.646372

**Authors:** Veronica Castle, Merdeka Miles, Rodrigo Fernandez-Gonzalez, Gonca Erdemci-Tandogan

**Affiliations:** Department of Cell and Systems Biology, University of Toronto, Toronto, ON, M5S 3G5, Canada; Translational Biology and Engineering Program, Ted Rogers Centre for Heart Research, University of Toronto, Toronto, ON, M5G 1M1, Canada; Department of Physics and Astronomy, University of Western Ontario, ON, N6A 3K7, Canada; Institute of Biomedical Engineering, University of Toronto, Toronto, ON, M5S 3G9, Canada; Developmental and Stem Cell Biology Program, The Hospital for Sick Children, Toronto, ON, M5G 1×8, Canada

## Abstract

Tissue boundaries pattern embryos, suppress tumours, and provide directional cues. Tissue boundaries are associated with supracellular cables formed by actin and the molecular motor non-muscle myosin II. Actomyosin cables generate tension that prevents cell mixing. Whether other cellular behaviours contribute to the formation of linear interfaces between cell populations remains unclear. In the *Drosophila* embryo, an actomyosin-based boundary separates the ectoderm from the mesectoderm, a group of neuronal and glial progenitors. Mathematical modelling predicted that cell divisions in the ectoderm challenge the mesectoderm-ectoderm (ME) boundary. Consistent with this, suppressing ectoderm cell divisions *in vivo* prevented cell mixing across the ME boundary when actomyosin-based tension was lost. Our mathematical model also predicted that cell divisions sharpen the ME boundary by reducing tension and increasing cell motility in the ectoderm. We found that inhibiting ectoderm divisions *in vivo* reduced boundary linearity. Using laser ablation and cell tracking, we demonstrated that cell divisions reduced junctional tension and increased cell movement in the ectoderm. Together, our results reveal that cell divisions facilitate cellular rearrangements to increase fluidity in a novel mechanism for boundary refinement.

## Introduction

Tissue boundaries are critical for patterning and growth during embryonic development (Meinhardt, 1983). In adults, tissue boundaries can minimize tumor malignancy by limiting cell invasion (Astin et al., 2010). Boundaries between tissues must withstand challenges from cell movement, cell division, or cell death (Monier et al., 2010), and thus, boundaries are often associated with the generation of mechanical force to maintain distinct cell populations apart (Heisenberg and Bellaïche, 2013). Despite their importance, the mechanisms by which boundaries are maintained and refined are still not well understood.

Tissue boundaries maintain cells populations apart to ensure proper tissue organization (Sharrock and Sanson, 2020; Wang and Dahmann, 2020). For example, in the *Drosophila* wing imaginal disc, two boundaries resist cell mixing due to proliferation: one boundary separates the anterior and posterior compartments of the disc (Sharrock and Sanson, 2020; Wang and Dahmann, 2020), and a second boundary separates the dorsal and ventral compartments (Garcia-Bellido et al., 1973; Morata and Lawrence, 1975). In the vertebrate hindbrain, boundaries between rhombomeres facilitate the differentiation of distinct regions (Lumsden and Krumlauf, 1996; Moens et al., 1998), and loss of rhombomere boundaries can cause abnormal development of the cranial and nasal bones (Twigg et al., 2004; Davy et al., 2006). Boundaries can also segregate cancerous cells from healthy tissues. In the mouse intestine, a boundary surrounds carcinomas, and metastasis can only occur if the boundary is disrupted (Cortina et al., 2007). Similarly, in the prostate, the boundary between epithelial and stromal compartments is usually lost during tumor invasion (Foty and Steinberg, 2004). Thus, boundaries are a key feature for the development and maintenance of normal tissue organization, and disruption of tissue boundaries is associated with disease (Major and Irvine, 2006; Landsberg et al., 2009; Monier et al., 2010; Calzolari et al., 2014; Yu et al., 2021).

Mechanical force is fundamental to the establishment and maintenance of tissue boundaries. Boundaries often display an enrichment of the molecular motor non-muscle myosin II and filamentous actin (F-actin), forming supracellular cables connected by cell-cell adhesive structures that span the length of the boundary (Wang et al., 2020). Actomyosin cables are present in the anterior-posterior and dorsal-ventral boundaries of the *Drosophila* wing disc (Major and Irvine, 2006; Landsberg et al., 2009; Umetsu et al., 2014), and also in the embryo, in the parasegmental boundaries that separate repeating developmental units (Monier et al., 2010), or around salivary gland precursors (Sanchez-Corrales et al., 2018). Beyond *Drosophila*, actomyosin cables characterize the boundaries between the notochord and presomitic mesoderm in *Xenopus* (Rohani et al., 2011) and around the eye field of zebrafish (Cavodeassi et al., 2013). Actomyosin cables at boundaries generate force that smoothens the interface between adjacent tissues (Landsberg et al., 2009; Monier et al., 2010; Calzolari et al., 2014).

The mesectoderm in the *Drosophila* embryo recently emerged as a system to study tissue boundaries (Yu et al., 2021). The mesectoderm separates the mesoderm (ventral) from the ectoderm (lateral) on both sides of the ventral midline (Video S1). At the end of mesoderm internalization, contralateral mesectoderm cells meet at the midline and seal the mesoderm inside the embryo. Mesectoderm cells then undergo a single round of oriented divisions that facilitate axis elongation (Wang et al., 2017; Camuglia et al., 2022). As the embryo develops, the mesectoderm is internalized, giving rise to glia and neurons of the central nervous system (Jacobs and Goodman, 1989; Klämbt et al., 1991; Tepass and Hartenstein, 1994; Wheeler et al., 2006).

A tissue boundary separates the mesectoderm from the ectoderm (Yu et al., 2021). After dividing, mesectoderm cells reverse their planar polarity and localize both actin and myosin II at the interface with the ectoderm, forming supracellular cables flanking the mesectoderm. The supracellular cables sustain increased tension. Increased tension at the ME boundary prevents ectoderm invasion of the mesectodermal domain, allowing the mesectoderm to internalize in a timely manner. Importantly, the actomyosin cable at the ME boundary is disassembled over time, in a process thought to contribute to the internalization of the mesectoderm.

Cell divisions challenge tissue boundaries. During division, cells round up through a reduction in adhesion to adjacent cells and an increase in osmotic pressure (Stewart et al., 2011; Fischer-Friedrich et al., 2014). Thus, dividing cells generate forces on adjacent cells. When cell divisions occur next to a boundary, they can transiently deform the boundary (Monier et al., 2010). Actomyosin contractility at the boundary generates tension in response to deformation, effectively pushing dividing cells into their original compartment and restoring the smooth interface between tissues. Notably, ectoderm cells divide next to the ME boundary (Wang et al., 2017). However, the impact of ectoderm cell divisions on the ME boundary has not been investigated.

## Results

### Mathematical modelling predicts that cell divisions challenge the ME boundary

We used computational modelling to investigate if ectoderm cell divisions play a role in the dynamics of the ME boundary. To this end, we developed an adaptive vertex model with time-varying parameters. The model was initialized with cell geometries corresponding to an *in vivo* configuration after mesectoderm cells divide and before ectoderm divisions begin (Fig. 1A, Video S2). Cells were assigned an energy that increased as cell area or perimeter deviated from a target value, with target values defined differently for ectoderm and mesectoderm cells based on *in vivo* measurements (Yu et al., 2021) (Materials and Methods). To simulate the myosin cables at the ME interface, we incorporated a time-dependent line tension between ectoderm and mesectoderm cells that decreased over time, consistent with the disassembly of the myosin cable at the ME boundary. Ectoderm cells were randomly selected to divide, with a frequency and orientation based on experimental data (Yu et al., 2021). Energy minimization guided the evolution of the cells in the model.

**Figure 1.**
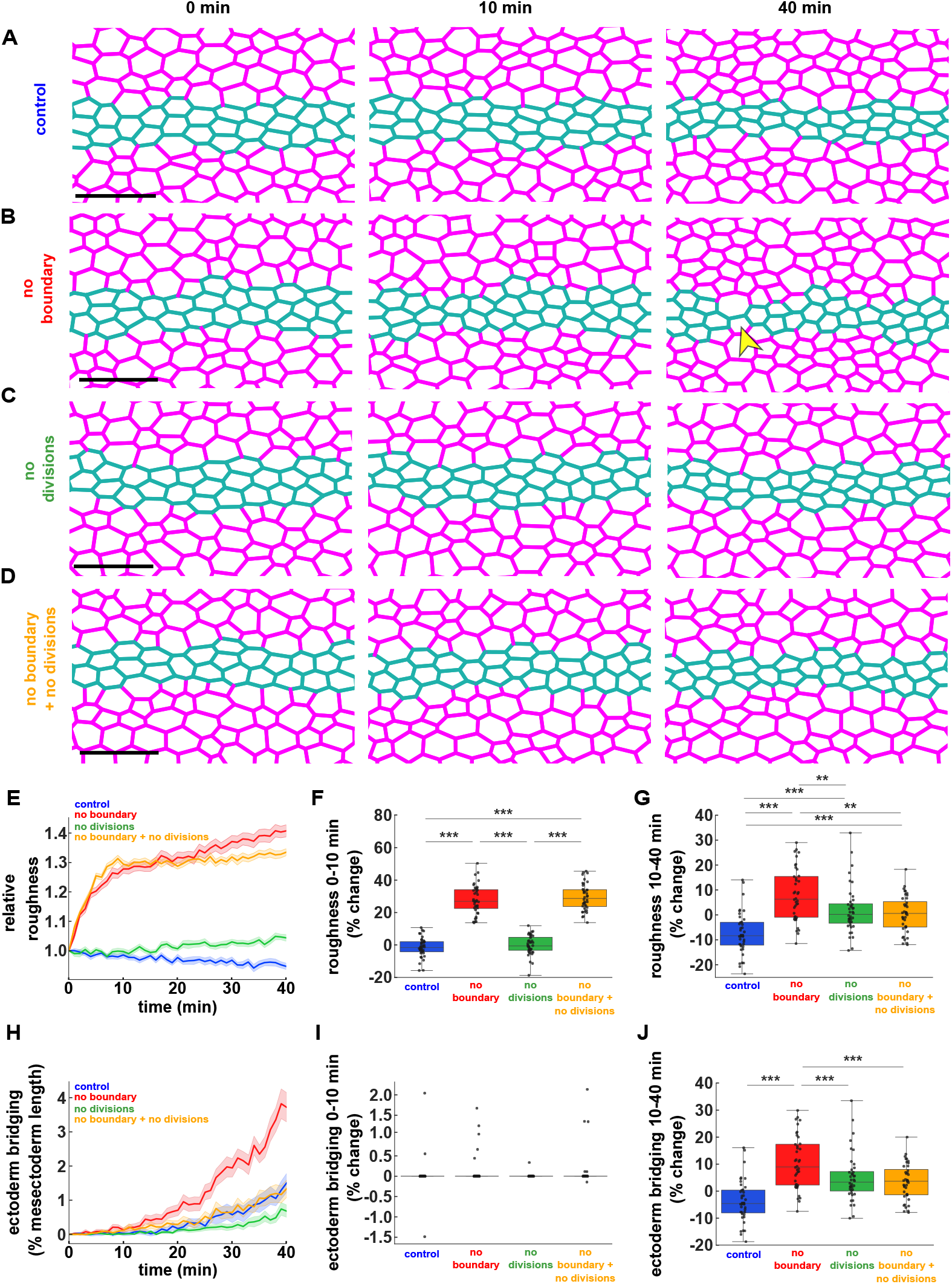
Mathematical modelling predicts that ectoderm divisions both challenge and refine the ME boundary. **(A-D)** Simulations of mesectoderm ingression in control embryos (A), with an acute loss of tension (no boundary) at the ME interface (B), without ectoderm cell divisions (C), or with simultaneous acute loss of tension at the ME interface and inhibition of ectoderm cell divisions (D). Magenta, ectoderm; teal, mesectoderm. Arrowheads indicate contralateral ectoderm cells in close proximity. Bars, 20 μm. Anterior, left. Time zero corresponds to the time in which the mesectoderm width starts decreasing. **(E-J)** Relative boundary roughness (E), change in boundary roughness from 0-10 minutes (F) or from 10-40 minutes (G), degree of ectoderm bridging along the AP axis (H), change in ectoderm bridging from 0-10 minutes (I) or from 10-40 minutes (J), in simulations of control embryos (blue, *n* = 40 simulations), embryos with acute loss of tension at the ME boundary (red, *n* = 40), embryos without ectoderm cell divisions (green, *n* = 40), and embryos with both acute loss of tension at the ME boundary and no ectoderm cell divisions (orange *n* = 40). (E, H) Error bars, s.e.m.. (F-G, I-J) Error bars, range; box, quartiles; grey lines, median. ** *P* < 0.01, *** *P* < 0.001.

Our computational model predicted that the ME boundary prevents cell mixing. We found that acutely reducing tension at the ME boundary significantly increased the roughness of the ME interface by 27±1% (mean±standard error of the mean, s.e.m.) within 10 minutes (*P* < 0.001), and by an additional 11±2% over the next 30 minutes, for a total 41±2% increase by 40 minutes (*P* < 0.001, Fig. 1A-B, E-G). Contralateral ectodermal cells came in close proximity or established premature contacts in the absence of ME boundaries (Fig. 1B, arrowheads). We defined premature contacts between contralateral ectoderm cells as ectoderm bridges, and we quantified the formation of ectoderm bridges by measuring the percentage of pixel columns along the anterior-posterior axis where the two ectoderm sheets were within 3 μm. We found that ectoderm bridges covered 3.7±0.4% of the mesectoderm length 40 minutes after removing the boundary, a significantly greater percentage than in controls (1.5±0.3%, *P* < 0.001, Fig. 1A-B, H-J). Thus, our model suggests that tension at the ME boundary maintains a smooth interface between ectoderm and mesectoderm and prevents cell mixing.

To investigate if the ME boundary prevents cell mixing by resisting ectoderm cell divisions, we used the model to quantify boundary dynamics when ectoderm divisions were inhibited (Fig. 1C-D). Inhibiting ectodermal divisions did not prevent the increase in ME boundary roughness 10 minutes after losing tension at the boundary, but reduced the secondary increase in roughness between 10 and 40 minutes by 68% (*P* < 0.01, Fig. 1E-G). Thus, modelling predicts that cell divisions provide a morphogenetic stress (not the only one) that challenges boundary linearity. Inhibiting ectodermal divisions in the model when the boundary was lost reduced the formation of ectoderm bridges by 77% at 40 minutes (*P* < 0.001 Fig. 1H-J), suggesting that ectoderm cell divisions promote cell mixing and are resisted by the ME boundary.

### Cell divisions challenge the ME boundary in living embryos

To test the prediction that the ME boundary resists ectoderm divisions to prevent cell mixing, we looked for methods to acutely inhibit ectoderm divisions *in vivo*, while still allowing mesectoderm cells to divide and the ME boundary to form. Dinaciclib inhibits cyclin-dependent kinases 1, 2, 5, and 9 (Parry et al., 2010), and it has been used to inhibit cell division in the *Drosophila* embryo (Akhmanova et al., 2022). We found that treatment with 500 μM of dinaciclib at the end of mesectoderm divisions prevented subsequent divisions in the ectoderm (Video S3). To disrupt the cable at the ME boundary, we treated embryos with 10 mM of Y-27632, a Rok inhibitor, as we did before (Yu et al., 2021) (Video S3). Notably, inhibiting myosin with Y-27632 did not prevent cell division (Video S4): we measured 1.0±0.2 completed cytokinesis/min in a 136×136 μm^2^ area of the ectoderm in control embryos, and 0.7±0.1 completed cytokinesis/min in embryos treated with Y-27632.

Similar to our previous findings (Yu et al., 2021), myosin inhibition *in vivo* disrupted the ME boundary. We quantified a rapid increase in boundary roughness, with an initial 20±5% increase within 10 minutes (*P* < 0.001), and a smaller, secondary increase of 6±2% over the next 30 minutes, for a total increase of 26±4% by 40 minutes (*P* < 0.001, Fig. 2A-B, E-G). Additionally, disrupting the myosin cable at the ME boundary led to the formation of ectoderm bridges that covered 16±6% of the mesectoderm length at 40 minutes (Yu et al., 2021) (*P* < 0.01, Fig. 2A-B, H-J). Thus, our results show that myosin activity is important to maintain a linear boundary and prevent cell mixing at the ME interface.

**Figure 2.**
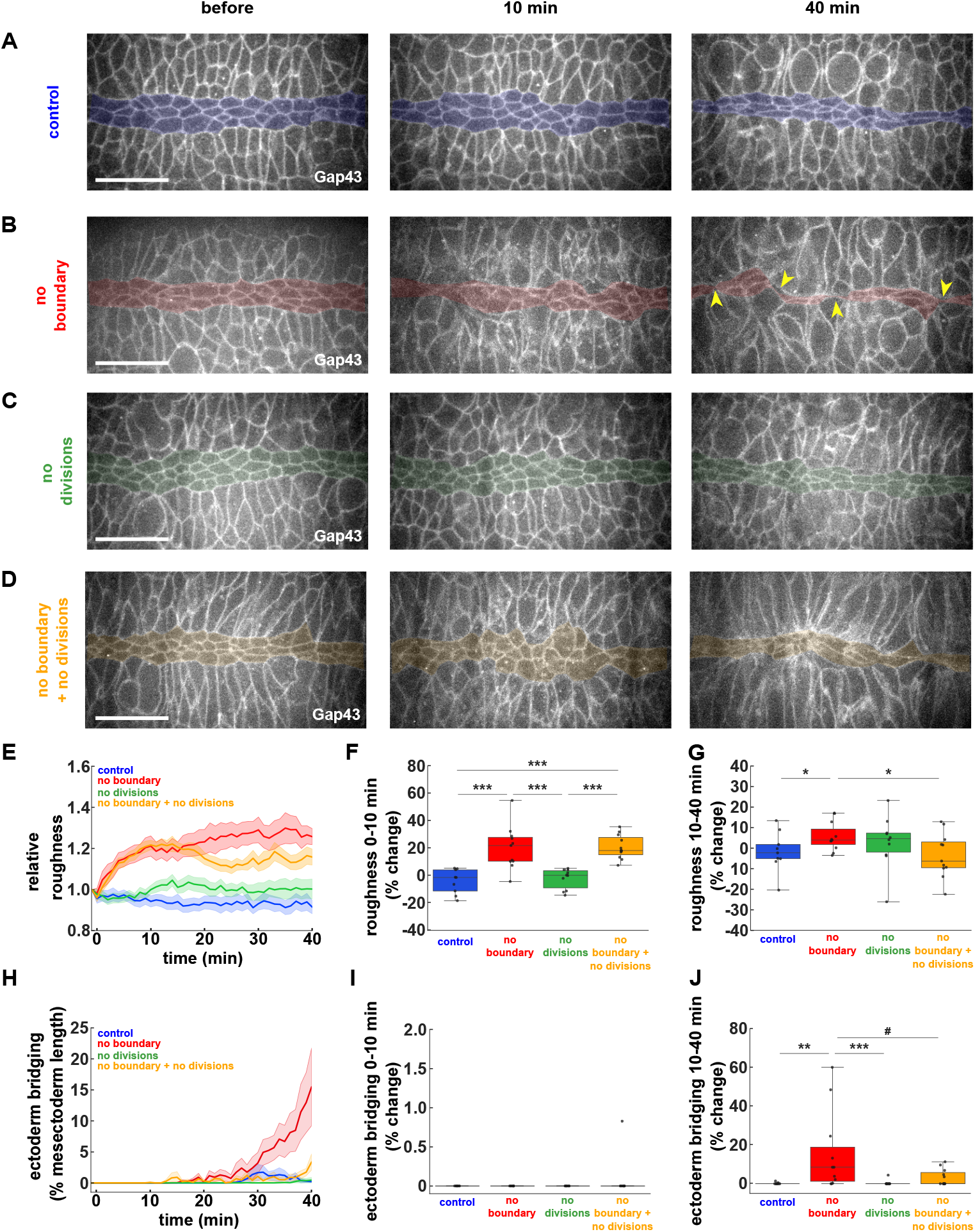
Ectoderm divisions both challenge and refine the ME boundary *in vivo*. **(A-D)** Mesectoderm (centre, shaded) and ectoderm (top and bottom) cells in embryos expressing Gap43:mCherry, and injected, 1 hour after the onset of mesectoderm divisions, with 50% DMSO (A), 20 mM Y-27632 (B), 500 μM dinaciclib (C), or both 20 mM Y-27632 and 500 μM dinaciclib (D). Arrowheads indicate contralateral ectoderm cells in close proximity. Bars, 20 μm. Anterior, left. **(E-J)** Relative boundary roughness (E), change in boundary roughness from 0-10 minutes (F) or from 10-40 minutes (G), degree of ectoderm bridging along the AP axis (H), change in ectoderm bridging from 0-10 minutes (I) or from 10-40 minutes (J), in embryos treated with 50% DMSO (blue, *n* = 9 embryos), 20 mM Y-27632 (red, *n* = 11), 500 μM dinaciclib (green, *n* = 11), or both 20 mM Y-27632 and 500 μM dinaciclib (yellow, *n* = 11). (A-H) Time is with respect to the time of injection. (E, H) Error bars, s.e.m.. (F-G, I-J) Error bars, range; box, quartiles; grey lines, median. # *P* = 0.06, * *P* < 0.05, ** *P* < 0.01, *** *P* < 0.001.

To determine if cell divisions deform the ME boundary when myosin is inhibited, we co-injected embryos with Y-27632 and dinaciclib after the ME boundary had formed, to simultaneously disrupt the ME boundary and prevent ectoderm cell divisions (Fig. 2A, D). Consistent with our *in silico* model predictions, inhibiting ectoderm divisions did not rescue the initial increase in boundary roughness associated with ME boundary disruption, but prevented the secondary increase in roughness at 40 min (*P* < 0.05, Fig. 2E-G) and reduced the formation of ectoderm bridges, which decreased to 3±1% of the mesectoderm length at 40 minutes (*P* < 0.06, Fig. 2J). Together our data indicate that ectoderm divisions *in vivo* challenge the ME boundary, suggesting that the ME boundary resists ectoderm divisions to prevent cell mixing.

### Cell divisions contribute to ME boundary linearity

Cell divisions typically challenge and deform tissue boundaries (Monier et al., 2010). Strikingly, our model predicted that cell divisions may contribute to the maintenance of the ME boundary. In control simulations, the boundary was refined slowly but continuously, with roughness decreasing by 5±1% within 40 minutes (Fig. 1A, E), consistent with a better-defined interface between ectoderm and mesectoderm. Suppressing cell divisions in the model reverted this trend, with roughness increasing (rather than decreasing) by 4±1% (Fig. 1C, E-G). Thus, mathematical modelling suggests that cell divisions in the ectoderm may contribute to the linearity of the ME boundary.

To test whether cell divisions refine the ME boundary *in vivo*, we quantified the roughness of the ME boundary in control embryos and in embryos treated with dinaciclib to inhibit cell division. Consistent with model predictions, we found that in control embryos, boundary roughness decreased significantly by 8±2% over 40 minutes (*P* < 0.05, Fig. 3A, C-D and Video S5). In contrast, inhibiting cell division prevented the reduction in ME boundary roughness (*P* < 0.05, Fig. 3A-B, C-D and Video S5). Together, our results show that cell divisions not only challenge, but also refine the tissue boundary between ectoderm and mesectoderm in the *Drosophila* embryo.

**Figure 3.**
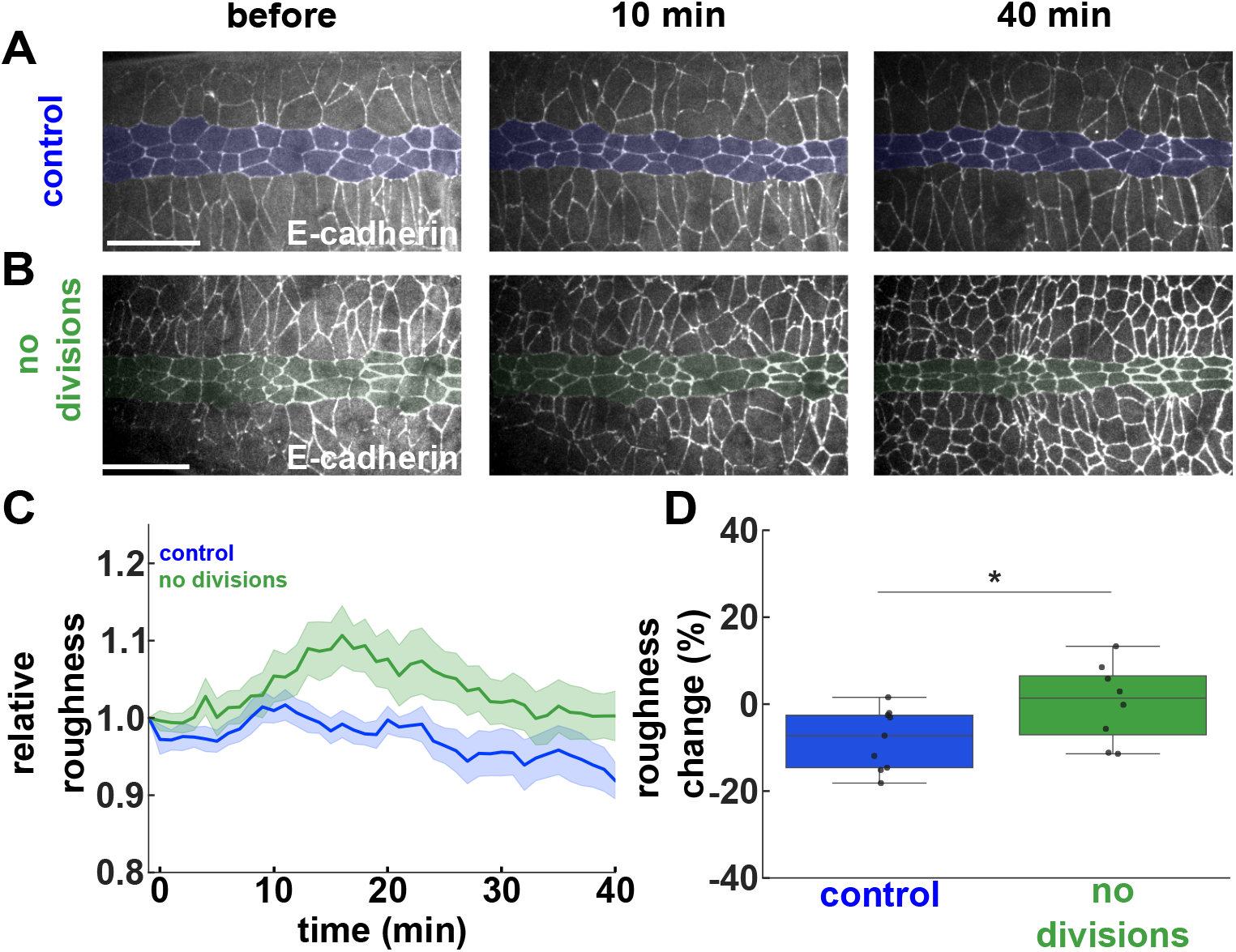
Ectoderm divisions increase the linearity of the ME boundary. **(A-B)** Mesectoderm (centre bracket) and ectoderm (top and bottom) cells expressing E-cadherin:GFP, and injected, 1 hour after the onset of mesectoderm divisions, with 50% DMSO (A) or 500 μM dinaciclib (B). Bars, 20 μm. Anterior, left. **(C-D)** Relative boundary roughness (C) and change in boundary roughness 40 min after injection (D) in embryos treated with 50% DMSO (blue, *n* = 9 embryos) or 500 μM dinaciclib (green, *n* = 9). (A-D) Time is with respect to the time of injection. (C) Error bars, s.e.m.. (D) Error bars, range; box, quartiles; grey lines, median. * *P* < 0.05.

### Cell divisions facilitate ectoderm cell movement

Cell divisions can fluidize tissues, facilitating the reorganization of cells (Lenne and Trivedi, 2022). To investigate if cell divisions refine the ME boundary by promoting the rearrangement of cells, we measured the mobility of ectoderm cells in the presence or absence of cell divisions (Fig. 4A-B). We measured cell mobility using a self-overlap function that characterizes glassy dynamics in molecular and colloidal materials (Castellani and Cavagna, 2005). Specifically, the self-overlap function measures the fraction of “static cells” or cells that moved a distance shorter than a characteristic length scale over a time period (Materials and Methods). In our computational model, the degree of self-overlap in the ectoderm in the presence of cell divisions decreased continuously, reaching half of its final value by 31±1 min (*P* < 0.001, Fig. 4A, C). In contrast, when we inhibited cell divisions in the model, the self-overlap function did not decrease for 78% of simulations (31/40), and for the remaining simulations the self-overlap only reached half of its final value by 38±1 min (Figure 4B-C), a significantly longer half-time than in the presence of cell divisions (*P* < 0.001), suggesting that the cells were frozen. We obtained similar results when we used the mean-squared displacement (MSD) of the cell centroids to quantify cell movements: in controls, the MSD of ectoderm cells at 40 min decreased by 83% when cell divisions were inhibited (*P* < 0.001, Fig. 4D). Thus, mathematical modelling predicts that cell divisions increase cellular mobility in the ectoderm.

**Figure 4.**
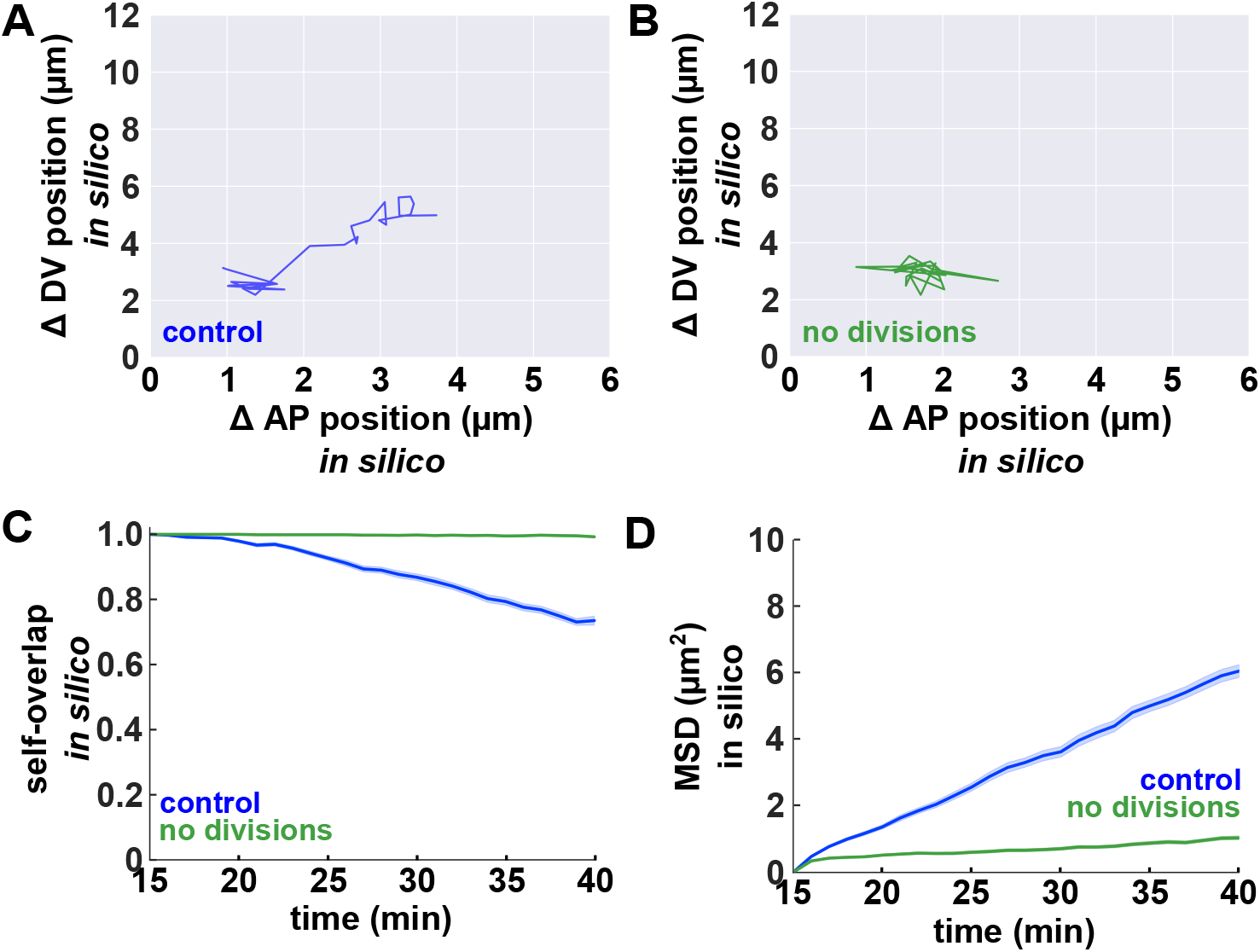
Mathematical modelling predicts that cell divisions increase ectoderm cell mobility. **(A-B)** Sample ectoderm cell centroid traces *in silico*, for controls (blue) or when cell divisions were inhibited (green). **(C-D)** Self-overlap function (C) and MSD (D) for ectoderm cells *in silico*, in the presence (blue) or absence (green) of cell divisions (*n* = 40 simulations per group, 40 cells per simulation). Error bars, s.e.m..

To test the prediction that cell divisions increase cellular mobility *in vivo*, we calculated the degree of self-overlap for ectoderm cells in control embryos or in embryos in which cell divisions had been blocked with dinaciclib (Fig. 5A-B). Following dinaciclib injection, cells that were already dividing completed their divisions. Thus, we began our analysis 15 minutes after injection. Consistent with our model predictions, ectoderm cells changed their positions rapidly in the presence of divisions, with a half-time of self-overlap decrease of 27±2 min (*P* < 0.001, Fig. 5A, C). In contrast, and as anticipated by the model, inhibiting cell divisions limited cell mobility in the ectoderm: the half-time of self-overlap decrease was 33±1 min, a significantly longer time than in controls (*P* < 0.05 Fig. 5B, C). Similarly, the MSD of the cell centroids decreased by 51% when we inhibited cell divisions (*P* < 0.01, Fig. 5D), further suggesting that *in vivo*, cell divisions fluidize the ectoderm. Of note, cell displacements when cell divisions were inhibited were greater *in vivo* that *in silico*, possibly due to additional forces, including intercalary cell behaviours in the ectoderm (Irvine and Wieschaus, 1994; Bertet et al., 2004; Blankenship et al., 2006), or the invagination of the mesoderm (Leptin and Grunewald, 1990; Sweeton et al., 1991; Clarke et al., 2025) and the posterior midgut (Collinet et al., 2015; Lye et al., 2015), that contribute to the movement of ectoderm cells in living embryos. Overall, our mathematical modelling and experimental results suggest that cell divisions may contribute to the refinement of the ME boundary by facilitating cellular movements.

**Figure 5.**
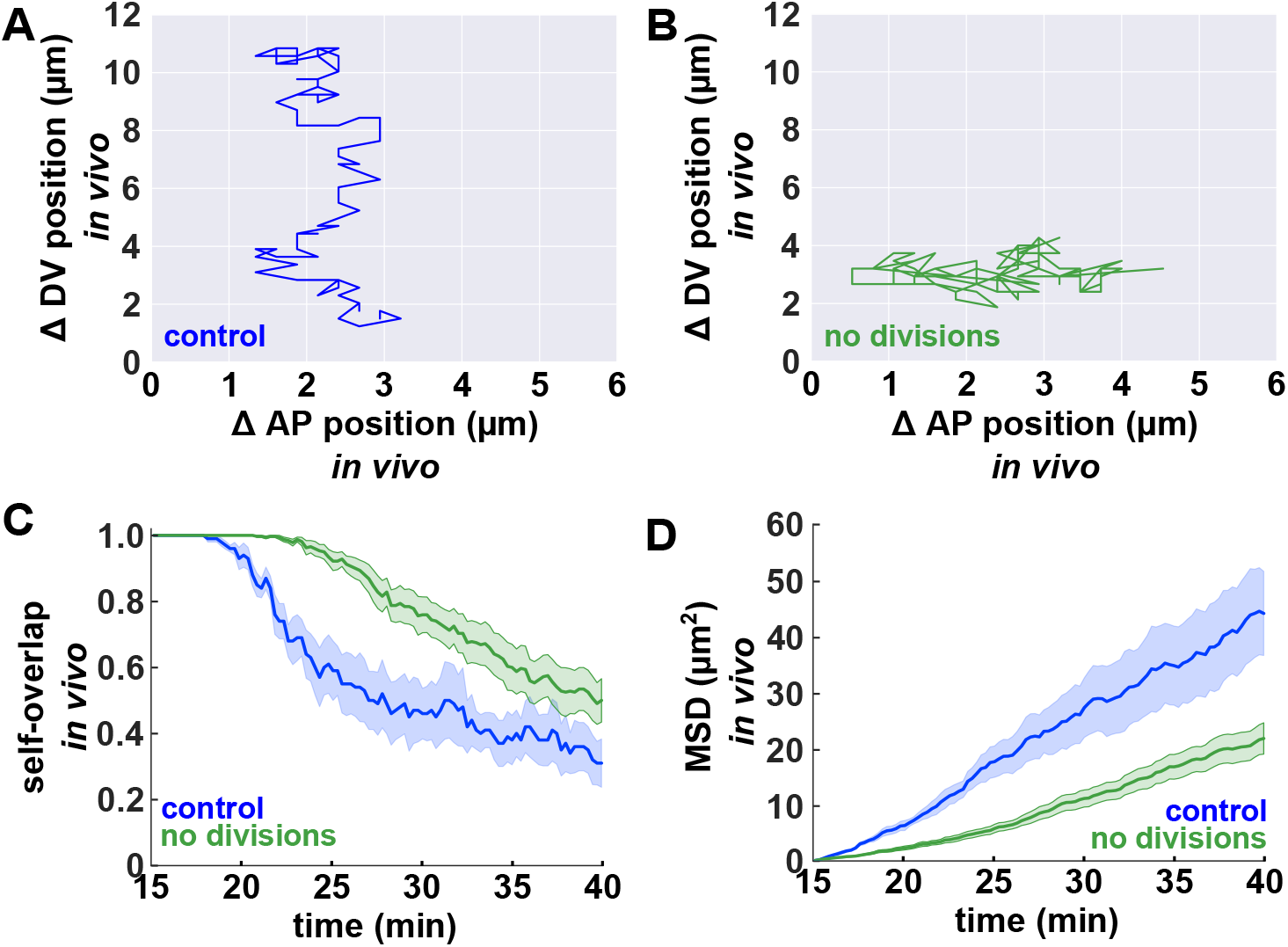
Cell divisions fluidize the ectoderm *in vivo*. **(A-B)** Sample ectoderm cell centroid traces *in silico*, for controls (blue) or when cell divisions were inhibited (green). **(C-D)** Self-overlap function (C) and MSD (D) for ectoderm cells *in vivo* in DMSO-treated controls (blue, *n* = 5 embryos, 20 cells per embryo) and in dinaciclib-treated embryos (green, *n* = 16 embryos, 20 cells per embryo). Error bars, s.e.m..

### Cell divisions release tension and increase fluidity in the ectoderm

Cell divisions can release tissue tension (Streichan et al., 2014; Wang et al., 2017). We speculated that preventing cell divisions may increase tension in the ectoderm, which would in turn limit cell mobility. To test this possibility, we first used our mathematical model to compare ectoderm tension in the presence and absence of cell divisions. We calculated junctional tension in the ectoderm directly from the energy of the cells (Materials and Methods) (Video S5). In the presence of cell divisions, tension at junctions between ectoderm cells decreased by 21.0±0.3% over the first 40 minutes of simulation (*P* < 0.001 Fig. 6A, C). Inhibiting cell divisions had the opposite effect on junctional tension, which increased by 5.5±0.2% (*P* < 0.01 Fig. 6B-C). Eventually junctional tension was 33% greater when cell divisions were inhibited than in controls (*P* < 0.001, Fig. 6C-D). Together, our modelling results predict that cell divisions release tension in the ectoderm.

**Figure 6.**
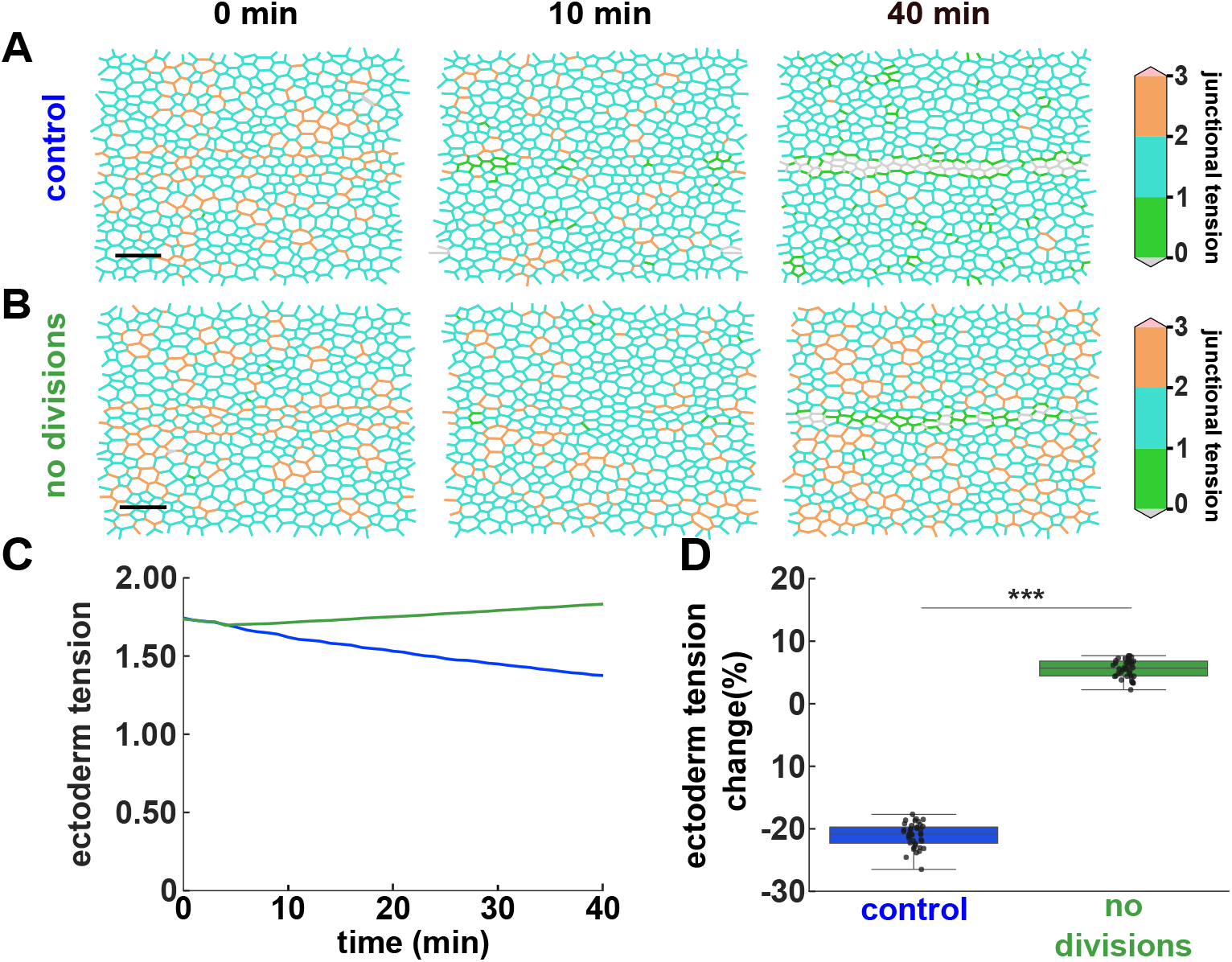
Mathematical modelling predicts that cell divisions reduce junctional tension. **(A-B)** Junctional tension distribution in simulations of mesectoderm ingression in controls (A) or when ectoderm cell divisions were inhibited (B). Bars, 20 μm. Anterior, left. **(C-D)** Junctional tension over time (C) and percent change in tension at 40 minutes (D) for ectoderm cells in control simulations (blue, *n* = 40 simulations, 1438 junctions on average per simulation) or in simulations with no ectoderm divisions (green, *n* = 40 simulations, 1233 junctions on average per simulation). (C) Error bars, s.e.m.. (D) Error bars, range; box, quartiles; grey lines, median. *** *P* < 0.001.

To test if cell divisions release ectoderm tension to facilitate cell remodelling *in vivo*, we used laser ablation to measure the tension sustained by individual cell-cell junctions in the ectoderm in both control and dinaciclib-treated embryos (Fig. 7A-B and Video S6). Under the assumption of uniform viscoelastic properties, the initial recoil velocity after ablation of a cell-cell junction is proportional to the tension that the junction sustained (Hutson et al., 2003; Zulueta-Coarasa and Fernandez-Gonzalez, 2015). We found that the recoil velocity after ablation of ectoderm cell junctions was 24% greater when cell divisions were inhibited than in controls (*P* < 0.05, Fig. 7A-C). Thus, our results indicate that cell divisions reduce tissue tension in the ectoderm during mesectoderm internalization.

**Figure 7.**
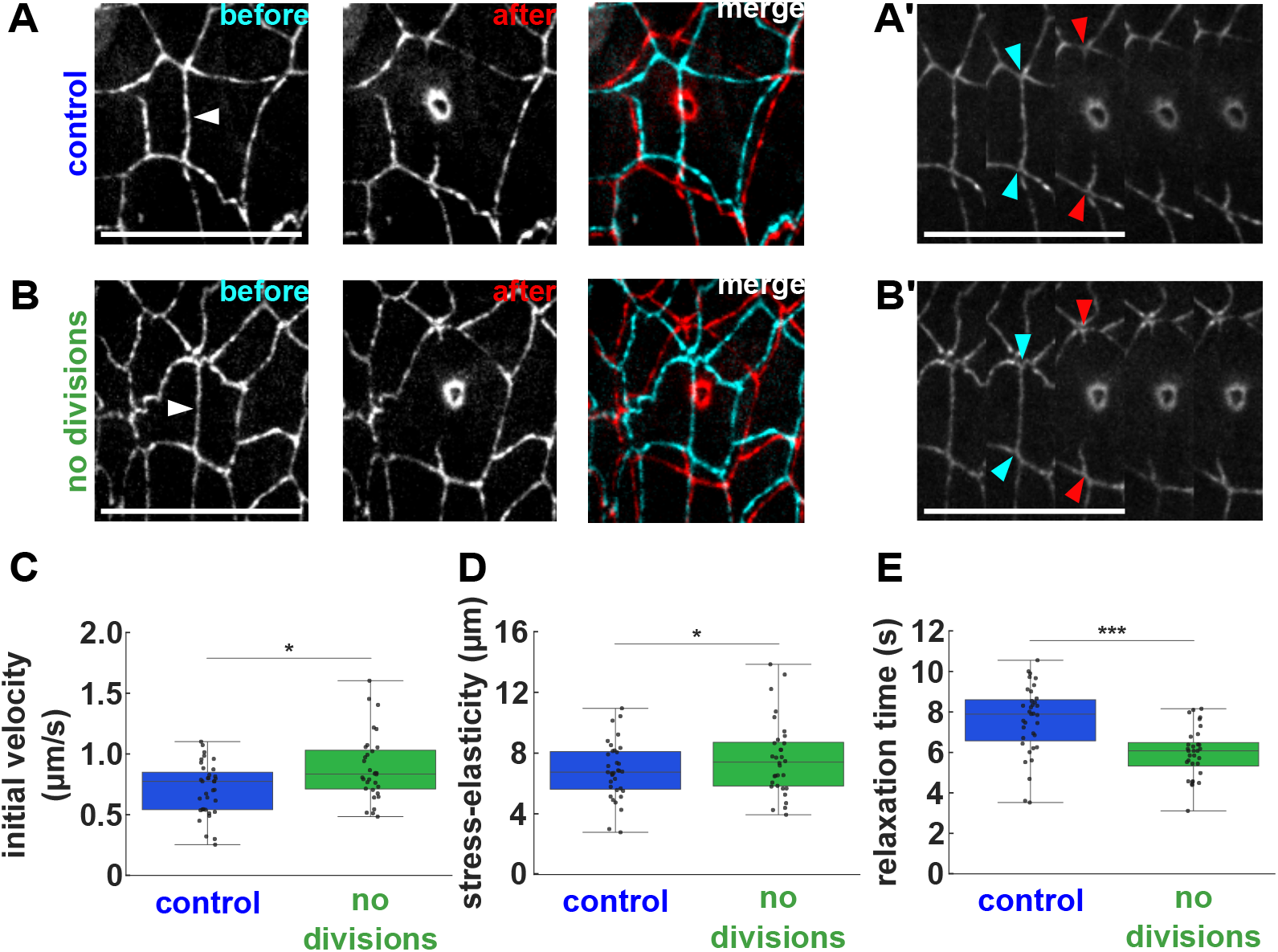
Cell divisions reduce junctional tension in the ectoderm *in vivo*. **(A-B)** Ectoderm cells expressing E-cadherin:GFP, immediately before (left, cyan in merge) and after (right, red in merge) ablation of a cell-cell junction parallel to the dorsal-ventral axis, in embryos treated with 50% DMSO (A) or 500 μM dinaciclib (B). Corresponding kymographs are shown (A’-B’). Arrowheads indicate the severed interface (white, A-B), or its ends prior to ablation (cyan, A’-B’) or immediately after (red, A’-B’). Bars, 20 μm (A-B) and 4 s (A’-E’). Anterior, left. **(C-E)** Initial recoil velocity after ablation (C), stress-elasticity ratio (D), and relaxation time (E) for cuts in embryos treated with 50% DMSO (blue, *n* = 35 cuts) or 500 μM dinaciclib (green, *n* = 32). (C) Error bars, s.e.m.. Error bars, range; box, quartiles; grey lines, median. * *P* < 0.05, *** *P* < 0.001.

To further test the effect of cell divisions on the fluidity of the ectoderm, we used a Kelvin-Voigt mechanical equivalent circuit to fit the laser ablation results. The Kelvin-Voigt model allowed us to estimate a relaxation time that indicates how long it takes for the laser-induced displacements to dissipate, as well as the stress-to-elasticity ratio (Materials and Methods). In agreement with our measurements of recoil velocity, the stress-to-elasticity ratio was 12% greater when cell divisions were inhibited (*P* < 0.05, Fig. 7D), further indicating that cell divisions reduce ectoderm tension. Additionally, we found that the relaxation time decreased by 27% when cell divisions were inhibited *(P* < 0.001 Fig. 7E), consistent with “freezing” of the tissue and the reduced cell mobility that we quantified. Together, our data indicate that cell divisions reduce junctional tension in the ectoderm, facilitating cell movement and tissue fluidity for ME boundary refinement.

## Discussion

Boundaries must withstand challenges from forces generated during development to ensure proper tissue patterning and cell fate specification. Using mathematical modelling, we predict that cell divisions within the ectoderm challenge the ME boundary. Surprisingly, our modelling also predicts that ectoderm divisions refine the ME boundary by releasing tension and fluidizing the tissue. *In vivo* experiments using pharmacological treatments, quantitative microscopy and laser ablation support modelling predictions, showing that ectoderm divisions play dual roles both challenging and refining the ME boundary as the embryo develops. Our results also show that adaptive vertex models with tissue-specific features from *in vivo* measurements and time-varying parameters are highly effective in capturing complex active processes during embryonic development (Tah et al., 2025).

The mechanisms that control myosin localization at the ME boundary are unclear. Tension is partly responsible for the polarized localization of myosin to the interface between ectoderm and mesectoderm (Yu et al., 2021), although the origin of that tensile force is not known. Posterior pulling forces, generated by the invagination of the posterior midgut could generate anisotropic stress along the anterior-posterior axis, dissipated by ectoderm cells through intercalation (Collinet et al., 2015; Lye et al., 2015), but not by mesectoderm cells, which do not exchange neighbours (Wang et al., 2017). This model would thus predict that mesectoderm cells would be the main contributors to the accumulation of myosin at the ME boundary, something that has not been tested. Beyond tension-based myosin localization, other mechanisms could drive the polarization of myosin at the ME boundary. The Eph/Ephrin receptor-ligand system promotes boundary formation. Eph/Ephrin signalling is associated with actomyosin accumulation at tissue borders (Cortina et al., 2007; Calzolari et al., 2014; Kindberg et al., 2021). In axons, Eph/Ephrin signalling activates the Rho GEF Ephexin via tyrosine phosphorylation, thus inducing Rho signalling and downstream actomyosin contractility (Sahin et al., 2005; Klein, 2012). Ephexin signalling also induces Rho signalling, actomyosin cable assembly and the formation of a boundary around damaged cells in developing embryos (Rothenberg et al., 2023). Importantly, overexpression of Ephrin in the mesectoderm causes defects in the ventral nerve cord (Bossing and Brand, 2002), suggesting that Eph/Ephrin signalling may contribute to proper mesectoderm development.

Our data indicate that cell divisions not only challenge, but also refine the ME boundary. Increased tension at compartment boundaries often results in smooth interfaces with reduced roughness (Landsberg et al., 2009; Monier et al., 2010; Calzolari et al., 2014). However, the ME boundary becomes smoother while myosin levels decrease (Yu et al., 2021). How is the refinement of the boundary accomplished? Theoretical studies suggest that the frequency of cell divisions influences the relaxation time of a viscoelastic tissue: tissues undergoing more divisions behaving in a more fluid manner (Ranft et al., 2010), as cell divisions disrupt the solid-like structure of the tissue. In gastrulating zebrafish embryos, oriented cell divisions within the plane of the enveloping layer alleviate tension and support tissue spreading (Campinho et al., 2013). Similarly, cell divisions in the mesectoderm release tension and facilitate axis elongation (Wang et al., 2017). We show that cell divisions reduce junctional tension in the ectoderm. Thus, by releasing tension and increasing ectoderm fluidity, cell divisions may enable cell movements that sharpen the ME boundary despite the loss of myosin.

The mechanisms by which cell divisions reduce tissue tension are unclear. In tissues with low levels of proliferation, cells can adopt a solid-like or jammed state as cell junctions mature (Garcia et al., 2015), making rearrangements less feasible (Lawson-Keister and Manning, 2021). Thus, cell divisions in the ectoderm may destabilize cell-cell junctions, thus preventing the build up of junctional actomyosin levels (Yu-Kemp et al., 2021) and tension. Studies quantifying how blocking cell divisions in the ectoderm affects the turnover of adherens junctions as well as cortical actomyosin levels may shed light on how cell division facilitates cell redistribution for boundary refinement.

In conclusion, our results suggest a dual role for ectoderm divisions on the ME boundary. Similar to other systems, divisions in the ectoderm challenge the ME boundary. Surprisingly, ectoderm divisions also refine the boundary. The mechanisms that establish and maintain tissue boundaries are conserved (Sánchez-Corrales and Röper, 2018). Thus, our findings may reveal a conserved mechanism whereby boundaries across proliferative tissues not only resist, but also benefit from cell divisions.

## Materials and Methods

### Fly stocks

We used the following markers for live imaging: *sqh*-*gap43:mCherry* (Martin et al., 2010), *sqh-sqh:GFP* (Royou et al., 2004), *sqh-sqh:mCherry* (Martin et al., 2009), *endo-e-cadherin:GFP* (Huang et al., 2009), *ubi-e-cadherin:GFP* (Oda and Tsukita, 2001).

### Time-lapse imaging

Stage 7-9 embryos were dechorionated in 50% bleach for 2 minutes, rinsed, glued ventral side down to a glass coverslip using heptane glue, and mounted in a 1:1 mix of halocarbon oil 27 and 700 (Sigma-Aldrich). Embryos were imaged using a Revolution XD spinning disc confocal microscope (Andor Technology) equipped with an iXon Ultra 897 camera (Andor Technology). The dynamics of mesectoderm internalization as well as the effects of drug treatments were imaged with a 60x oil immersion lens (Olympus, NA 1.35). Sixteen-bit Z-stacks were acquired at 0.5 μm steps every 4-60s (15-27 slices per stack), and maximum intensity projections were used for analysis.

### Drug injections

Embryos were dechorionated and glued to a coverslip as above, dehydrated for 7.5 minutes, and covered with a 1:1 mix of halocarbon oil 27 and 700. Embryos were injected using an M-LSM motorized micromanipulator (Zaber), and a PV820 microinjector (WPI) attached to a spinning disk confocal microscope. Drugs (Y-27632, Tocris Bioscience, and dinaciclib, ApexBio) were injected into the perivitelline space where they are predicted to be diluted 50-fold (Foe and Alberts, 1983). Y-27632 was injected at 10 mM in 50% DMSO, and dinaciclib was injected at 500 μM in 50% DMSO; control embryos were injected with 50% DMSO. Drugs were injected 1 hour after the first mesectoderm division.

### Laser ablation

Ablations were induced using a Pulsed Micropoint N_2_ laser (Andor) tuned to 365nm. The laser delivers 120 μJ pulses of 2-6 ns each. Ten pulses were delivered at a single point to sever cell-cell junctions in the ectoderm. Samples were imaged immediately before and every 4 s after ablation. Laser cuts were conducted 30 minutes following drug treatment.

To estimate changes in viscoelasticity, we modelled cell-cell contacts as viscoelastic elements using a Kelvin-Voigt mechanical-equivalent circuit (Zulueta-Coarasa and Fernandez-Gonzalez, 2015). The Kelvin-Voigt circuit represents junctions as the combination of a spring (elasticity) and a dashpot (viscosity) configured in parallel. Considering the equations that represent the forces sustained by a spring and a dashpot, it is possible to derive the equation for the change in length between the ends of the junction after ablation:

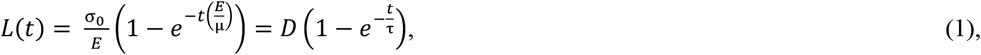

where *L*(*t*) is the distance between the ends of the ablated junction at time *t* after ablation, *σ*_*0*_ is the tension sustained by the junction, *E* is the elastic coefficient, and *μ* is the viscosity. Using the laser ablation data it is possible to estimate the asymptotic value of *L, D*, and a relaxation time, *τ*, that estimates the viscosity-to-elasticity ratio.

### Image segmentation and analysis

Image analysis was performed using our open-source image analysis platforms, PyJAMAS (Fernandez-Gonzalez et al., 2021) and SIESTA (Fernandez-Gonzalez and Zallen, 2011; Leung and Fernandez-Gonzalez, 2015). To segment mesectoderm boundaries, we used the LiveWire algorithm in PyJAMAS, an interactive method based on Dijkstra’s minimal cost path algorithm (Dijkstra, 1959) to find the brightest pixel path between any two pixels in an image. Cell boundaries were segmented using a combination of the LiveWire algorithm and the active contour method *balloons*, implemented in PyJAMAS, in which a polygon evolves on an image towards its minimum energy configuration, with the polygon energy inversely related to the image gradient, and with a balloon force that ensures polygon expansion over regions of the image with small image gradients (Zulueta-Coarasa et al., 2014).

To measure the linearity of the ME boundary, we rotated images so that the anterior-posterior axis of the embryo was oriented along the horizontal axis of the image, and we quantified the roughness of the boundary, *ω*_*L*2_ (Michel et al., 2016), defined as:

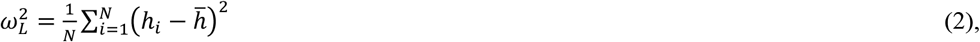

where *N* is the number of pixel columns along a 13-μm segment, *L*, of the ME boundary, *h*_*i*_ is the *y* coordinate the *i*-th pixel in the segment, and 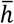 is the average *y* coordinate of the pixels in the segment. Thus, *ω*_*L*2_ represents the local variance of the position of the ME boundary with respect to the anterior-posterior axis of the embryo.

To measure the formation of ectoderm bridges at time *t*, we quantified *R*, the fraction of pixel columns along the anterior-posterior axis where the two ectoderm sheets were within 3 μm:

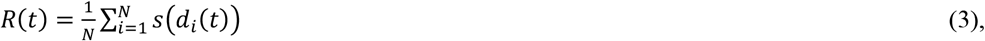

with *d*_*i*_(*t*) the distance between opposite ectoderm sheets on the *i*-th pixel column at time *t*, and *s* the step-function:

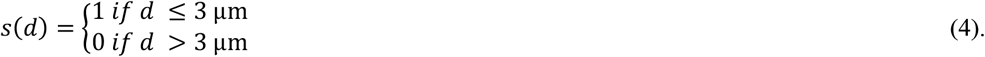

To measure cell movement in the ectoderm, we quantified a self-overlap function, *Q* (Castellani and Cavagna, 2005), representing the fraction of cells at time *t* that moved by less than 4 μm (approximately one cell radius (Yu et al., 2021)):

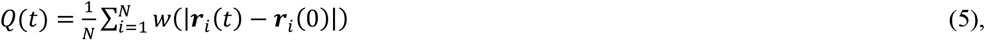

where ***r***_*i*_ is the position of *i*-th cell centroid, and *w* is a step function that weighs the change in cell centroid position:

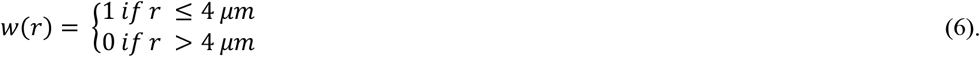

We validated our self-overlap results using the mean squared displacement of ectoderm cell centroids, defined as:

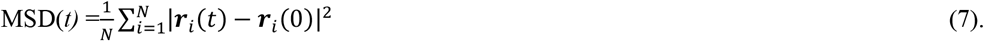

Only cells ectoderm cells that had not divided were used for the self-overlap and MSD analyses.

### Vertex model

We used an adaptive 2D vertex model with cell divisions and time-varying parameters based on *in vivo* measurements. Our implementation utilized the open-source framework cellGPU (Sussman, 2017). In vertex models, cells are depicted as collections of nodes (vertices) and edges (interfaces between cells), which represent a cross-section of the tissue (Yu and Fernandez-Gonzalez, 2017). The energy, *E*, of the tissue is determined based on the geometry of the cells:

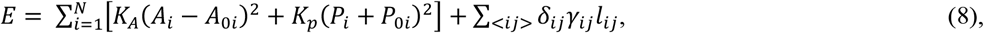

where *N* denotes the number of cells in the tissue. The first term of Eq. 7 is the incompressibility of the cell: *K*_A_ is an area spring constant and *A*_*i*_ and *A*_*0i*_ are the current and preferred areas of the *i*-th cell, respectively. The second term represents the competition between adhesion and contractility: *K*_P_ is a perimeter spring constant, and *P*_*i*_ and *P*_*0i*_ are the current and preferred perimeters of the *i*-th cell, respectively. The final term represents the myosin cable between mesectoderm and ectoderm cells: *δ*_*ij*_ is 1 when the adjacent cells *i* and *j* belong to different tissue types, otherwise *δ*_*ij*_ is zero; *γ*_*ij*_ represents the tension generated by the myosin cable at the interface between ectoderm and mesectoderm cells; and *l*_*ij*_ denotes the length of the contact between cells *i* and *j*.

The evolution of the model is guided by an energy minimization process. We used the forward Euler method to update the position of each cell vertex:

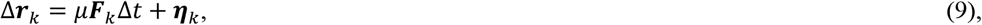

where ***r***_*k*_ is the position of vertex *k*, ***F***_*k*_ = − ∇_*k*_*E* is the force on vertex *k, μ* is the inverse friction coefficient, Δ*t* is the integration time step, and ***η***_*k*_ is a normally distributed random force with zero mean and variance 2*μT*Δ*t*. The temperature *T* allows the system to have Brownian fluctuations in the position of cell vertices (Brańka and Heyes, 1999). The natural unit length of the simulations is given by 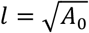. We set the integration time step Δ*t* = 0.01*τ* where *τ* = 1/(*μK*_*A*_*A*_*0*_) is the natural time unit of the simulations.

We initialized the simulations with 400 random cells in a periodic box. All simulations were run at their target parameters for 10^5^τ before setting ectoderm and mesectoderm cells features. We simulated the system after the completion of cell divisions in the mesectoderm and the formation of the myosin cables at the ME boundary. Thus, we set a common preferred area, *A*_0*mesec*_, for all mesectoderm cells (Yu et al., 2021) (Table S1). The preferred cell area for mesectoderm cells,

*A*_0*mesec*_, was updated over time based on experimental data (Yu et al., 2021). We modelled the ectoderm as a bidisperse mixture of cells representing before and after division, with preferred areas, 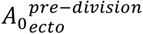 and 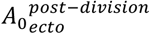, respectively (Yu et al., 2021) (Table S1). We scaled the preferred perimeter, *P*_*0*_, for all cell types to maintain a constant target shape index defined as 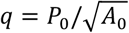 (Bi et al., 2015). We modelled the tension reduction at the ME boundary cell-cell interfaces using an exponential decay function 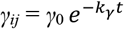, where *γ*_0_ is a constant line tension representing the myosin cable at the boundary at *t* = 0, and *k*_*γ*_ is the rate of tension reduction. We select *k*_*γ*_ values based on experimental myosin measurements at the boundary in control embryos and in embryos in which myosin activity was inhibited (Yu et al., 2021) (Table S1). Ectoderm cells were randomly chosen to divide, with frequency and cell division orientation determined experimentally (Yu et al., 2021). Following each division, the system was allowed to relax to a new energy minimum. To model the impact of inhibiting the cell divisions, ectoderm cell divisions were blocked at 5 minutes in the simulations to mimic the delayed effect of dinaciclib injection. Other parameter values (Table S1) were selected to minimize the difference between *in vivo* and *in silico* measurements of ME boundary roughness and ectoderm bridging. We assigned the natural time unit of the simulation to be *τ* = 0.1 min.

We calculated junctional tension in the model directly from the energy of the cells ((Yang et al., 2017):

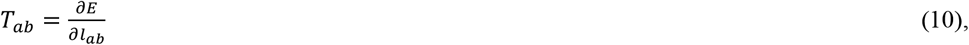

where *T*_*ab*_ is the tension along the edge that connects vertices *a* and *b*. Using Eq. (7), junctional tension in the ectoderm becomes:

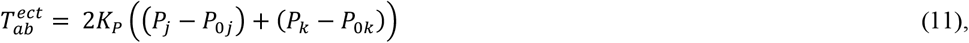

where *a* and *b* are the vertices that separate cells *j* and *k*.

### Statistical analysis

For multiple group comparisons, we used a Kruskal-Wallis test to reject the null hypothesis, followed by Dunn’s test for pairwise comparisons (Glantz, 2002). Dividing and non-dividing cells were compared using a non-parametric Mann-Whitney test, or Wilcoxon signed-rank test for paired data. For time series, error bars indicate the s.e.m. For box plots, error bars show the range, the box indicates the quartiles, and grey lines denote the median.

## Supplementary video legends

**Video S1. The mesectoderm separates the mesoderm (ventral) from the ectoderm (lateral) on both sides of the ventral midline**. Mesectoderm and ectoderm cells in a *Drosophila* embryo expressing E-cadherin:GFP. Mesectoderm cells are highlighted in blue. Time is with respect to the completion of mesectoderm divisions. Anterior, left. Images were acquired every 5 minutes.

**Video S2. Mathematical modelling predicts that ectoderm divisions both challenge and refine the ME boundary**. Simulations of mesectoderm ingression in control embryos (top left), with an acute loss of tension at the ME interface (bottom left), without ectoderm cell divisions (top right), or with simultaneous acute loss of tension at the ME interface and inhibition of ectoderm cell divisions (bottom right). Magenta, ectoderm; teal, mesectoderm. Anterior, left. Time zero corresponds to the time in which the mesectoderm width starts decreasing. Images were generated every 10 minutes.

**Video S3. Dinaciclib treatment inhibits ectoderm divisions and prevents ME boundary refinement**. Mesectoderm (centre) and ectoderm (top and bottom) cells in embryos expressing E-cadherin:GFP, and injected, 1 hour after the onset of mesectoderm divisions, with 50% DMSO (left) or 500 μM dinaciclib (right). Anterior, left. Images were acquired every 30 seconds.

**Video S4. Y-27632 treatment does not inhibit ectoderm divisions**. Mesectoderm (centre) and ectoderm (top and bottom) cells in embryos expressing myosin:GFP (green) and Gap43:mCherry (magenta), and injected, 1 hour after the onset of mesectoderm divisions, with 50% DMSO (left) or 20 mM Y-27632 (right). Anterior, left. Images were acquired every 30 seconds.

**Video S5. Mathematical modelling predicts that cell divisions reduce junctional tension in the ectoderm**. Junctional tension distribution in simulations of mesectoderm ingression in controls (left) or when ectoderm cell divisions were inhibited (right). Anterior left. Time zero corresponds to the time in which the mesectoderm width starts decreasing. Images were generated every 10 minutes.

**Video S6. Cell divisions reduce junctional tension in the ectoderm**. Laser ablation of a contact between ectoderm cells and parallel to the dorsal ventral axis in embryos expressing E-cadherin:GFP and treated with 50% DMSO (left) or 500 μM dinaciclib (right). Images were acquired every 4 seconds.

## Supplementary tables

**Table S1.** Simulation parameters in natural units.

|  |  |
| --- | --- |
| area spring constant, $K_A$ | 1 |
| perimeter spring constant, $K_P$ | 1 |
| inverse friction coefficient, $\mu$ | 1 |
| temperature, $T$ | 0.01 |
| initial preferred area of mesectoderm cells, $A_{0mesec}^{initial}$ | 0.79 |
| preferred area of ectoderm cells before division, $A_{0ecto}^{pre-division}$ | 1.32 |
| preferred area of ectoderm cells after division, $A_{0ecto}^{post-division}$ | 0.66 |
| target shape index of mesectoderm cells, $q_{mesec}$ | 3.4 |
| target shape index of ectoderm cells, $q_{ecto}$ | 3.4 |
| cell division time, $t_{cd}$ | 2 |
| tension constant, $\gamma_0$ | 0.4 |
| rate of acute tension reduction, $k_{\gamma}^{no\ boundary}$ | 0.36 |
| rate of tension reduction representing control embryos, $k_{\gamma}^{control}$ | 0.01 |

